# The novel phenomenon of inter-retinal coupling predicts cortical neurovascular responses in health and disease

**DOI:** 10.1101/2025.10.23.684136

**Authors:** João Jordão, Pedro Guimarães, Pedro Serranho, João Figueira, Miguel Morgado, Delia Cabrera DeBuc, Michel Paques, Miguel Castelo-Branco, Rui Bernardes

## Abstract

It has recently been discovered that physiological crosstalk exists between human retinas. Accordingly, visual stimulation of one retina elicited a neurovascular coupling (NVC) response of the contralateral retina in about 30% of individuals. The functional significance of this feature in health and disease remains unknown. Here, we asked the question whether inter-retina crosstalk predicts the nature of neurovascular coupling in the brain. To test this hypothesis, we assessed BOLD fMRI (blood-oxygen-level-dependent functional magnetic resonance imaging signal) in healthy individuals and a disease model of neurovascular coupling, type 1 diabetes. We used three visual paradigms: a Threshold speed and Sub-maximum speed discrimination tasks, and flicker stimulation. The presence or absence of inter-retinal coupling was predictive of the time course of the cortical hemodynamic response, either during the initial or late phases. The presence of interocular coupling predicted larger BOLD responses. Diabetic patients with interocular coupling showed a very similar profile to the healthy group. In sum, we identified a direct relationship between interocular physiological coupling and cortical hemodynamic responses, suggesting that this NVC response may serve as a disease biomarker in diabetes.

**Graphical Abstract:** 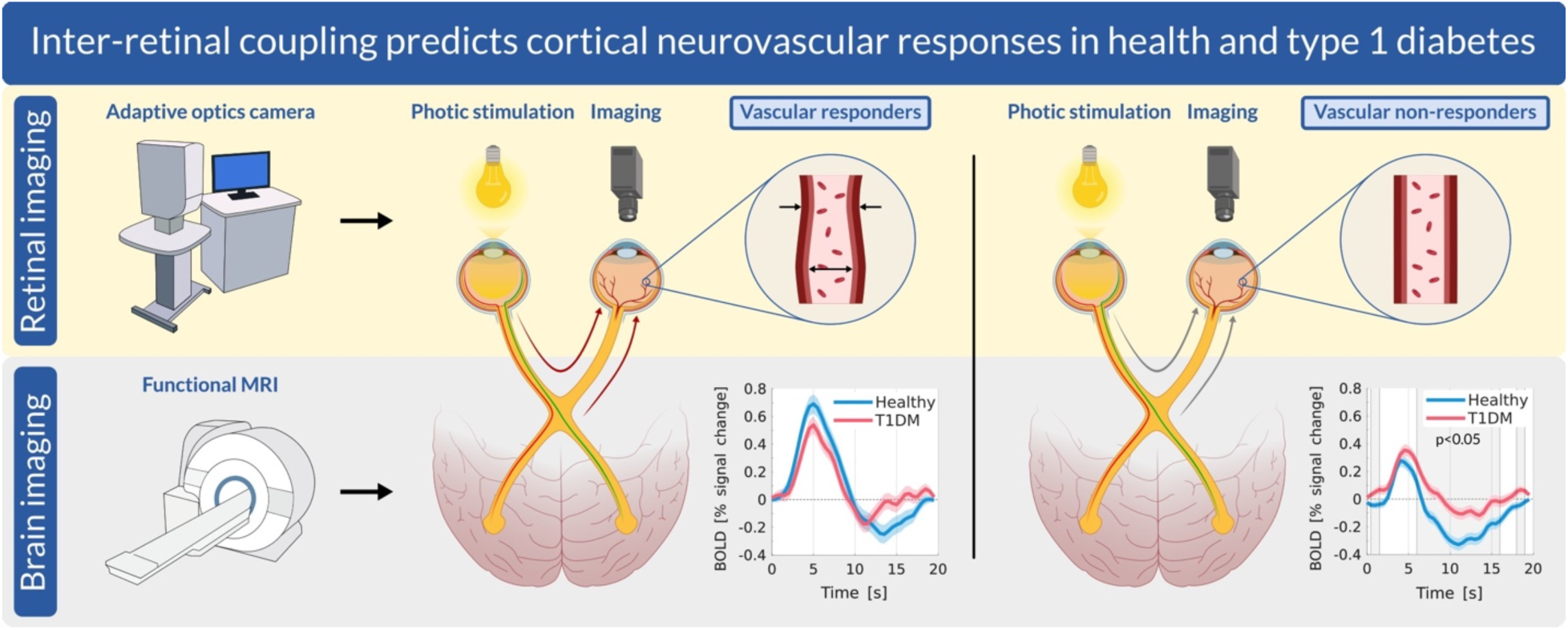

## 1. Introduction

Blood flow in the central nervous system (CNS) is tightly coupled to neuronal activity through a process known as neurovascular coupling (NVC), which ensures adequate delivery of oxygen and glucose in response to increased metabolic demand (Iadecola, 2017). The foundations of NVC lie in the lack of energy reservoir in the CNS, despite being a highly demanding system when stimulated (Willie et al., 2014; Iadecola, 2017). Indeed, blood flow must reach the CNS at the right place and in due time, as failure has severe consequences (Iadecola, 2017). Mechanisms of regulation have been extensively studied because of sheer scientific curiosity about how the brain regulates its blood supply and the realization of the harmful effects of its dysregulation. While it is clear that NVC is a key component of the healthy brain, whether its deficit drives pathology or the other way around is still largely unknown (Stackhouse and Mishra, 2021).

NVC dysfunction is accepted to be not only associated with Alzheimer’s disease (AD) (Iadecola, 2017), but also with several other disorders, including stroke (Phillips et al., 2016) and diabetes (Garhöfer et al., 2020; Canna et al., 2022; Duarte et al., 2023; Guimarães et al., 2024). Although the last decades’ research shed light on several cellular mechanisms underlying NVC, these are not yet fully characterized (Stackhouse and Mishra, 2021). Establishing the connection between the NVC for distant regions within the CNS may provide further insights into the complex NVC control system and its behavior at the CNS full scale, rather than focusing only on local effects.

We have recently discovered (Jordão et al., 2025) that at least one functional interocular crosstalk mechanism exists across adult human retinas. We demonstrated that photic stimulation of one retina elicits a vascular response in the contralateral (non-stimulated) retina. The fact that the stimulation of one region of the CNS may impact far-off regions may be of wide significance. This crosstalk mechanism was found in over 1/3 of individuals regardless of their health status (presence or absence of type 1 diabetes). Interocular neurovascular responses were measured by the variation of the lumen of retinal vessels in the macular region imaged by a high-resolution adaptive-optics fundus camera (Jordão et al., 2023; Jordão et al., 2025). Current models of NVC, such as those proposed by Girouard and Iadecola (2006), do not account for the recently identified interocular crosstalk mechanism, suggesting the need to revise or expand existing frameworks to incorporate these novel pathways. Here, we asked the question whether this new phenomenon is associated with changes in cortical hemodynamic responses as measured by the BOLD signal (blood-oxygen-level-dependent signal) using fMRI (functional magnetic resonance imaging). Long-range effects across the CNS remain poorly understood and require further attention (Sirotin and Das, 2009). These long-range effects were experimentally demonstrated by our group (Jordão et al., 2025) and include positive and negative neurovascular responses (Jordão et al., 2025). Negative neurovascular responses are prominent during early development and have been observed in both human infants and rodent models (Stackhouse and Mishra, 2021). In the adult brain, such responses are also evident in regions surrounding active visual areas (Bressler et al., 2007), though it remains unclear whether they arise from blood flow diversion toward more active regions or from local neural suppression. These observations raise the possibility that distinct hemodynamic response patterns may co-exist within the brain, reflecting a complex interplay between excitation, inhibition, and vascular dynamics (Hill et al., 2021).

BOLD-fMRI enables the indirect measurement of neuronal activity by capturing hemodynamic signals associated with NVC, which are directly related to the concentration of oxygen in red blood cells. The BOLD signal allows for the identification of brain regions activated in response to a given stimulus and the characterization of individual hemodynamic responses (Arichi et al., 2012). The hemodynamic response function (HRF) describes vascular responses to neuronal activation over time and is characterized by several key components: a) an initial dip, where the BOLD signal initially drops to a negative value; b) a positive response (peak); and c) an undershoot before returning to the baseline, see Fig. 1.

**Fig. 1.**
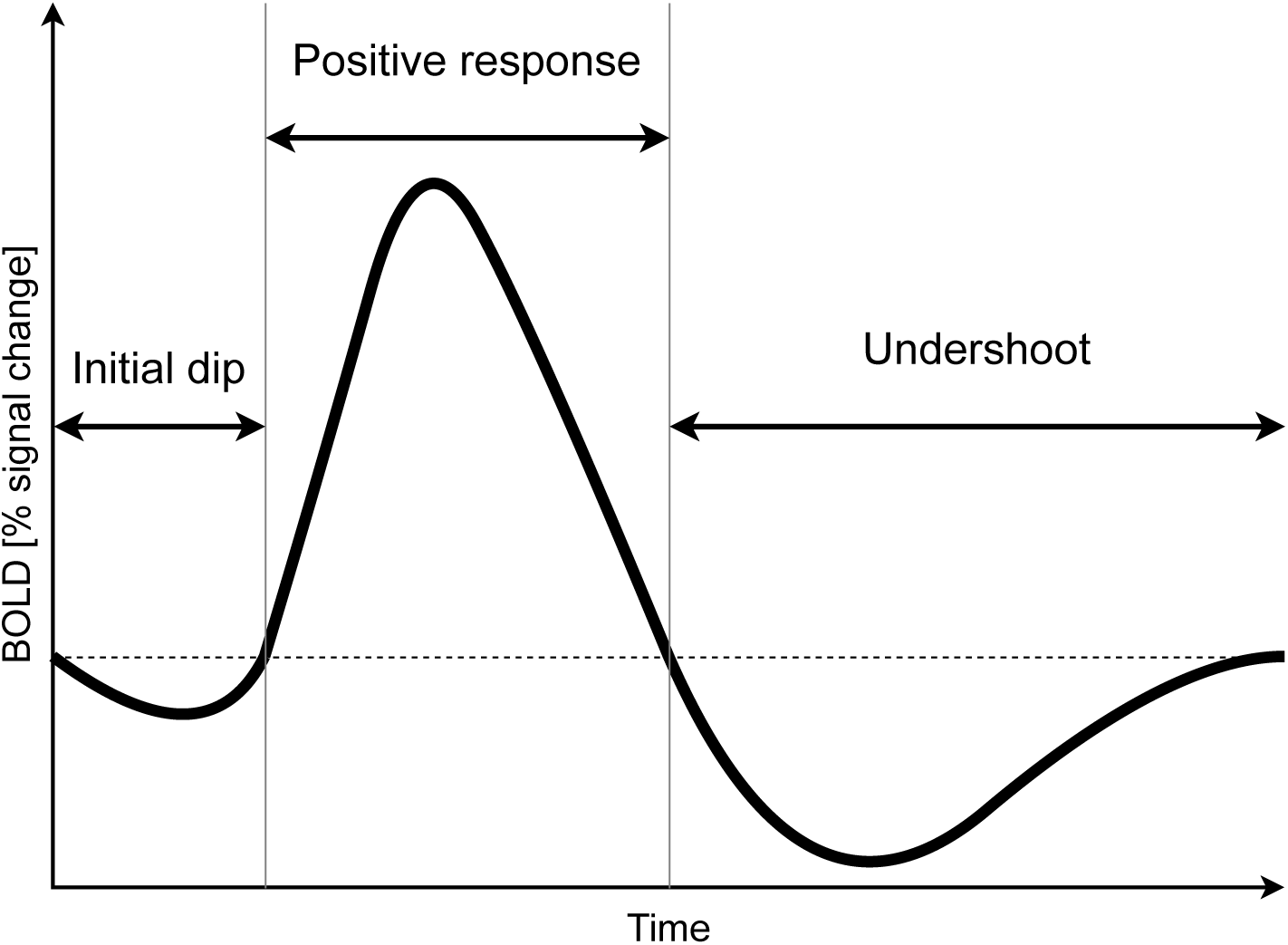
Simplified representation of the canonical hemodynamic response function. Key components: a) initial dip after stimulus onset; b) positive response period (comprising the peak); c) undershoot region after the peak and before recovering to the baseline.

Accurate staging of type 1 diabetes mellitus (T1DM) has become increasingly important for both clinical management and research, as it enables earlier identification of disease onset, more precise patient stratification, and tailored therapeutic interventions. Recent classification frameworks divide T1DM into distinct stages, ranging from asymptomatic autoimmunity (Stage 1) to dysglycemia (Stage 2) and overt hyperglycemia (Stage 3), providing a more nuanced understanding of disease progression and risk (Insel et al., 2015). This stratified approach is essential for identifying early biomarkers of disease, such as NVC alterations, which may precede clinical symptoms and offer insights into subclinical CNS involvement in T1DM. In this study, we provide a detailed comparison of the hemodynamic response between healthy and T1DM subjects as well as within two subgroups of each group. We translate our findings on the characterization of NVC differences in highlighting the need to better discriminate subjects enrolled in BOLD-fMRI studies according to their subtypes and evaluate the potential use of the retinal vascular network response to the contralateral retina photic stimulation to assess NVC onset changes associated with T1DM.

## 2. Research Design and Methods

### 2.1. Study design

The primary objective of this work was to evaluate the feasibility of utilizing a safer, faster, and cost-effective alternative imaging technique based on the study of NVC responses in the contralateral retinal vascular network and to compare it with the brain hemodynamic responses for the characterization of NVC onset changes associated with T1DM. Initially, the subjects underwent fMRI scanning using both block and event-related designs. Functional imaging acquisition runs comprised speed discrimination tasks and flicker stimuli patterns for the computation of brain hemodynamic responses. Subsequently, the subjects underwent retinal imaging using an adaptive optics fundus camera to measure vascular network changes to flicker stimulation of the non-imaged eye.

### 2.2. Recruitment

Data from 32 healthy subjects and 20 T1DM patients, with no signs of diabetic retinopathy, collected between October 2023 and July 2024, with all subjects fulfilling a set of inclusion and exclusion criteria disclosed in (Jordão et al., 2025), were accessed. A subgroup of 16 healthy controls (HC) (9 females, 7 males; age range: 19–55 years; median: 24.5 years; mean ± SD: 33.8 ± 12.8 years), along with all 20 T1DM patients (10 females, 10 males; age range: 20–57 years; median: 32.0 years; mean ± SD: 35.0 ± 12.6 years), underwent comprehensive ophthalmological examinations. These participants, 16 HC and 20 T1DM, had their retinal NVC responses evaluated using high-resolution adaptive optics fundus imaging and successfully completed their fMRI sessions to assess brain NVC responses to multiple visual stimuli.

For healthy controls, systolic blood pressure (BP) ranged from 98 to 130 mmHg (mean ± SD: 114.4 ± 10.0 mmHg), and diastolic BP ranged from 59 to 88 mmHg (mean ± SD: 70.9 ± 8.0 mmHg). In the T1DM group, systolic BP ranged from 90 to 139 mmHg (mean ± SD: 117.0 ± 13.3 mmHg), and diastolic BP ranged from 60 to 93 mmHg (mean ± SD: 73.8 ± 8.1 mmHg). Glycated hemoglobin (HbA1c) levels in the T1DM group ranged from 5.8% to 7.9%, with a median of 6.8% and a mean ± SD of 6.9 ± 0.5%. Detailed demographic information can be found in (Jordão et al., 2025) (Table 1). Selected subjects’ IDs are: HC13-17, HC19-22, HC25-26, HC28-32; and DM01-20 (Jordão et al., 2025). All participants provided written informed consent before participation. The study was conducted following the ethical principles of the Declaration of Helsinki and its subsequent revisions. Additionally, ethical approval (CE-039/2023) was obtained from the Ethics Committee of the Faculty of Medicine of the University of Coimbra.

**Table 1.**
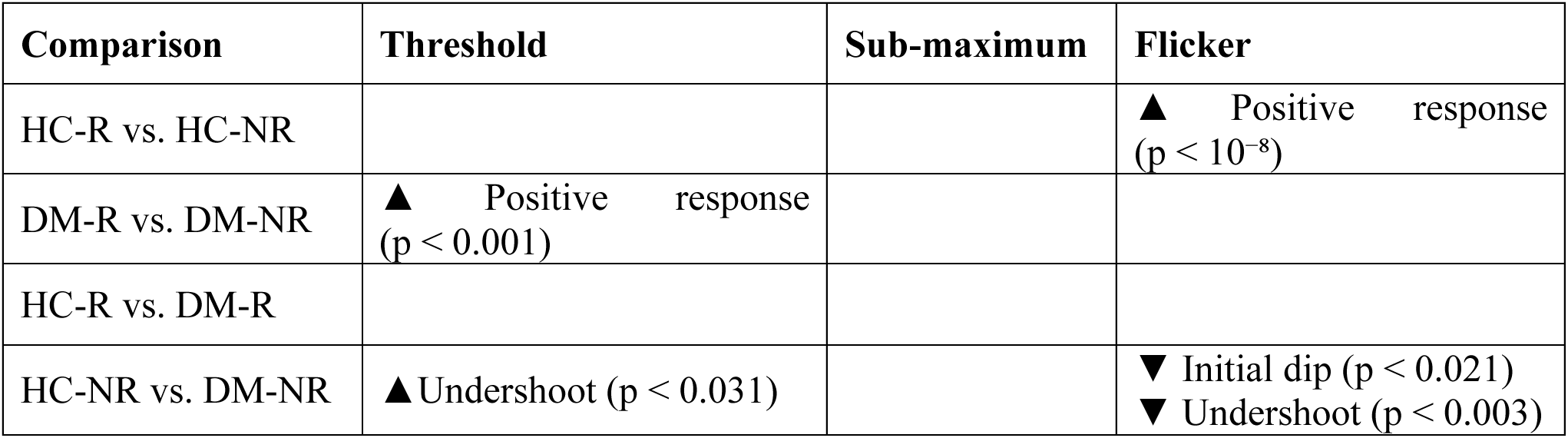
Simplified results of the healthy and T1DM subgroups’ hemodynamic response curve analysis. The matrix representation herein follows the one presented in Fig. 5. Only the comparisons with statistically significant p-values (p < 0.05) are shown, with their respective minimum reached p-value. The HC-NR vs. DM-NR comparison presents both initial dip and undershoot regions separately. The ▲ arrow indicates that the first subgroup has a higher BOLD response relative to the second, while the ▾ arrow indicates otherwise.

### 2.3. Brain imaging setup

All fMRI acquisitions were performed using the Siemens Magnetom 3T XR Numaris/X VA30A-03GR (Siemens, Germany) and comprised a 3D brain anatomical magnetization-prepared rapid acquisition gradient echo (MPRAGE) configured with a repetition time (TR) of 2530 ms, an echo time (TE) of 3.5 ms, an inversion time (TI) of 1100 ms, a flip angle of 7°, 192 slices, a voxel size of 1 mm × 1 mm × 1 mm, and a field-of-view (FOV) of 256 px × 256 px (256 mm × 256 mm); and four echo planar imaging (EPI) functional runs configured with a TR of 1000 ms, a TE of 37 ms, a flip angle of 68°, 66 slices, a voxel size of 2 mm × 2 mm × 2 mm, and an FOV of 110 px × 110 px (220 mm × 220 mm). Subjects viewed functional runs’ visual stimuli projected onto a 485 cm × 878 cm area on a screen positioned at a distance of 1749.7 cm (FOV of 15.41° vertical x 28.17° horizontal) through a mirror placed at 45°. Subjects responded to stimulus sequences using one control box (Lumina LS-Pair, Cedrus, CA, USA). No ophthalmic drops or contrast agents were administered during the procedure.

### 2.4. fMRI sequence and stimuli

The fMRI sequence, presented in Fig. 2, comprised a psychophysics task to assess individual speed discrimination threshold, a brain anatomical acquisition, one block-design run with a speed discrimination task, one block-design run with flicker stimuli, one event-related design run with a speed discrimination task, and one event-related run with flicker stimuli. The procedure remained identical for all participants, with the psychophysics task lasting an average of 78 seconds, and the remaining acquisitions lasting 28 minutes and 52 seconds.

**Fig. 2.**
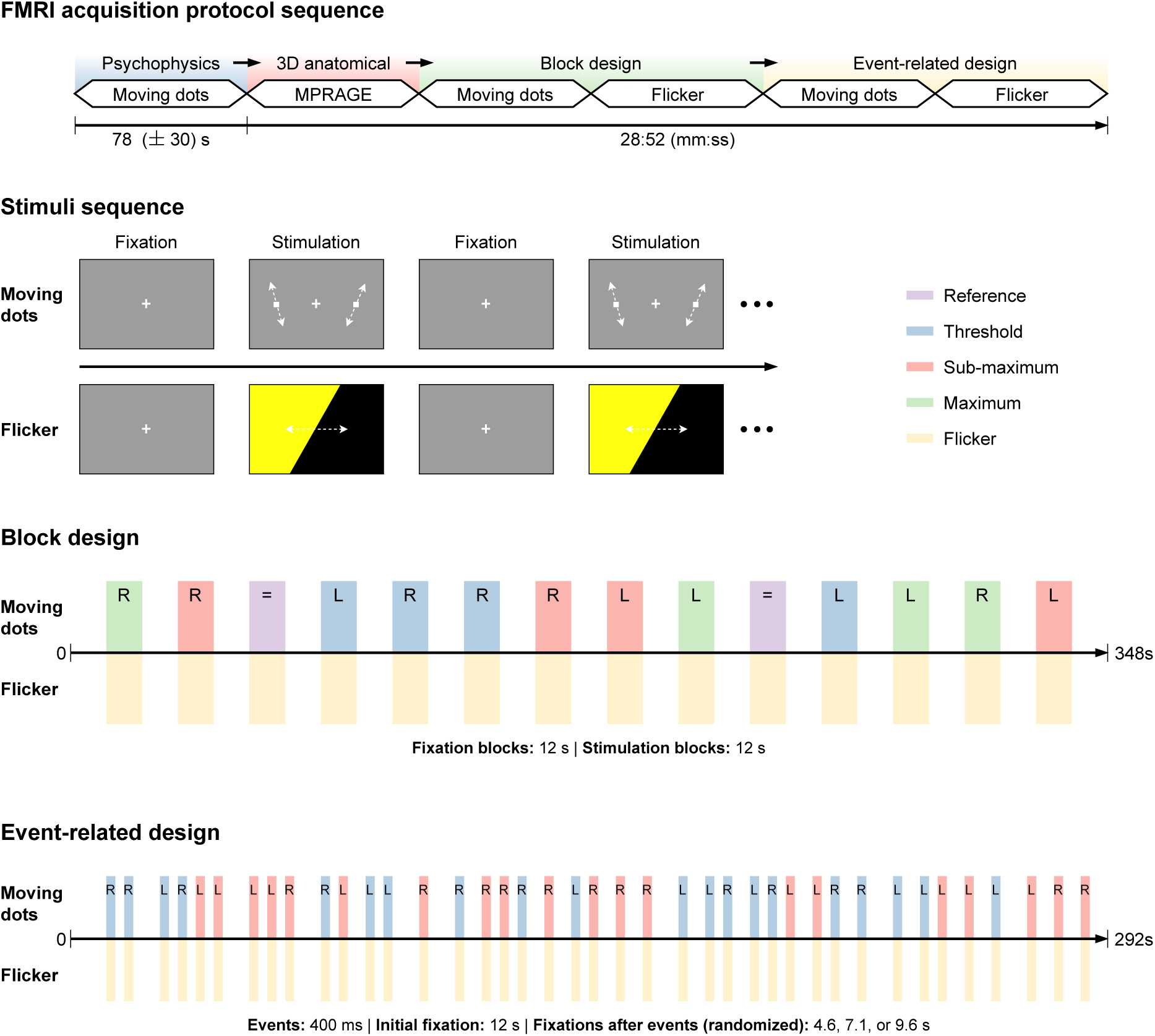
Graphical representation of the fMRI acquisition protocol sequence, stimuli, and experimental protocols of block and event-related designs. Top: Representation of the fMRI acquisition protocol sequence with configured duration. Stimuli sequence: Representation of the stimuli presented to the subjects. The fixation periods presented a white fixation target at the center of the screen. The moving dots stimulation representation shows examples of allowed dot movement directions. The flicker stimulation representation shows the flickering from a full yellow to full black screen. Block design: Diagram of the sequence of stimulation blocks discriminated by type. Event-related design: Diagram of the sequence of stimulation events discriminated by type.

The speed discrimination task showed two linear moving white dots, one on each vertical half of a dark gray background. The flicker stimuli covered the entire viewing screen, periodically changing between yellow (RGB triplet: 48.974 cd/m², 196.644 cd/m², 0.323 cd/m²) and black (RGB triplet: 0.148 cd/m², 0.137 cd/m², 0.150 cd/m²) at a frequency of 15 Hz (see Fig. 2).

### 2.5. Psychophysical task

Participants first completed a psychophysical task to determine their individual speed discrimination thresholds by identifying the faster of two moving dots, ensuring equivalent performance conditions across subjects. One dot, the reference dot, always moved with a constant speed of 5°/s, while the other, the faster-moving dot, started moving with a speed of 24°/s. The stimulus (the two dots moving) was presented for 400 ms, and the location (left or right) of the faster-moving dot relative to the reference dot was randomly selected each time these were shown to participants. The speed of the faster-moving dot was adjusted to decrease following a correct answer and increase otherwise. This continued until participants completed six reversals in response patterns (i.e., a correct answer followed by an incorrect one, or vice versa). The individual speed threshold was determined as the average speed at which the last four reversals occurred. Details can be found in (Duarte et al., 2015).

### 2.6. Block-design runs

The block-design runs consisted of 15 fixation periods without stimulation interleaved with stimulation periods for a total of 29 periods (blocks), each lasting 12 s (see Fig. 2). The moving dots stimulation blocks comprised one dot always moving with a reference speed of 5°/s and one dot moving with one of four allowed speeds: the reference speed of 5°/s (Reference), the reference speed plus the individual threshold (Threshold), the reference speed plus three times the threshold (Sub-maximum), and an arbitrarily established speed of 20°/s (Maximum). The Reference condition was presented twice, while the remaining conditions (Threshold, Sub-maximum, and Maximum) were presented four times each, twice with the faster-moving dot on the left side of the screen and twice on the right. The flicker block-design runs presented the 15 Hz yellow flicker (squared wave pulses) in all 14 stimuli blocks.

### 2.7. Event-related design runs

Each event-related run began with an initial fixation period of 12 s, followed by 40 events, each lasting 400 ms (see Fig. 2). This was followed by a fixation period of either 4.6 s, 7.1 s, or 9.6 s, randomly determined to minimize anticipation effects that might confound event-related responses (Friston et al., 1999). The entire sequence was established before the experiment and remained consistent across all subjects, thereby maintaining consistent response conditions across event-related runs. The moving-dots stimuli for the event-related sequence used the Threshold and Sub-maximum conditions defined above, thereby leaving out the Reference and Maximum conditions. Each condition was presented 20 times, with the faster-moving dot appearing 10 times on each side of the screen. The flicker events presented the stimulus 40 times.

### 2.8. Retinal imaging

Following fMRI acquisitions, one eye of each subject was imaged using the rtx1 adaptive optics fundus camera (Imagine Eyes, Orsay, France). Eyes were selected alternately between the right and left eyes, per group, at the study enrollment. Retinal images of 4° × 4° field-of-view, 1500 px ×1500 px, were obtained at an artery presenting a well-defined lumen and walls, and as close as possible to the subject’s fovea. All acquisitions of the same subject were performed at the same location. To assess NVC in the retinal vascular network, photic stimulation was employed using the following sequence of three images per condition: imaging without stimulation (Basal 1), imaging following the stimulation of the non-imaged eye (Contralateral), imaging without stimulation (Basal 2), and imaging following the stimulation of the imaged eye (Ipsilateral). Full details can be found in (Jordão et al., 2025).

### 2.9. Hemodynamic response in the retina

Arterioles were manually segmented using custom software to determine the average lumen width for a segment in prior work (Jordão et al., 2025). These can be found in Table S1 (supplementary material). The contralateral and ipsilateral NVC responses were computed as the percentage of the relative variation from the stimulus to the basal width. The contralateral responses were then evaluated using two criteria to assess individual relevant responses:

> C1: Only cases where the lumen’s width following a contralateral stimulation was either both over basal1 and basal2 lumens (positive response) or under basal1 and basal2 lumens (negative response).

> C2: The contralateral response should be over 1% or exceed 50% of the respective ipsilateral response.

Data for all cases are presented in Fig. S1 (supplementary material), and data that fulfill criteria C1 and C2 are shown in Fig. S2 (supplementary material).

### 2.10. Consensual retinal vascular response subgroups

Two subgroups were defined within both the healthy control and T1DM subject populations based on the responsiveness of their retinal vascular networks to contralateral photic stimulation. Subjects with a detectable retinal vascular response were classified as responsive (HC-R, n = 5 for healthy controls; DM-R, n = 7 for T1DM), while those without a measurable response were categorized as non-responsive (HC-NR, n = 11; DM-NR, n = 13). BOLD signals were analyzed separately for each of the four subgroups, and intergroup comparisons were performed to assess differences in neurovascular coupling dynamics.

### 2.11. fMRI data pre-processing

The fMRI data were pre-processed using BrainVoyager version 22.4 (Brain Innovation, Maastricht, Netherlands) (Goebel et al., 2006) and MATLAB version R2024b (The MathWorks Inc., Natick, Massachusetts). Anatomical MPRAGE scans, block-design runs, and event-related runs underwent distinct pre-processing pipelines. Anatomical volumes were skull-stripped and corrected for intensity inhomogeneities in BrainVoyager, isolating the brain and brainstem.

The block-design functional volumes underwent multiple steps: a) 3D motion correction using trilinear interpolation during realignment and sinc interpolation for the final transformation; b) slice timing correction using cubic spline interpolation, informed by the slice acquisition timing from the scan protocol; c) spatial smoothing using a Gaussian kernel with a full width at half maximum (FWHM) of 4 mm; and d) temporal filtering using a high-pass filter with a cutoff frequency of 0.015 Hz (Smith et al., 1999) to remove low-frequency drifts.

Functional volumes from the event-related design underwent the same motion correction procedure as the block-design data. Slice scan time correction was then performed in MATLAB because stimuli do not aligned with the start of the acquisition (TR of 1000 ms at present and 2500 ms in (Duarte et al., 2023)) in consequence of the use of random fixation periods defined in (Duarte et al., 2023), with advantages stated in (Friston et al., 1999), and BrainVoyager cannot handle these non-synchronous stimuli (SOA - stimulus onset asynchrony). Slice scan time correction was performed using cubic spline interpolation and evaluated at a temporal resolution of 500 ms because BrainVoyager can only handle ISI (inter-stimulus interval) multiples of TR. Temporally corrected volumes were imported back into BrainVoyager to proceed with temporal filtering using the same high-pass filter applied in the block-design runs, with a cutoff frequency of 0.015 Hz. Unlike the block-design data, event-related volumes were not spatially smoothed.

Finally, all functional volumes were co-registered to each participant’s anatomical scan using boundary-based registration and normalized to Montreal Neurological Institute (MNI) space.

### 2.12. Computation of brain neurovascular activation maps

The pre-processed block-design data from all participants were analyzed using a standard random-effects (RFX) general linear model (GLM). This approach estimates the percentage change in the BOLD signal for each voxel, allowing the results to be generalized to the broader population (Penny and Holmes, 2007). The BOLD signal brain maps from all moving dot conditions were combined and evaluated as a single entity. The resulting BOLD signals for the moving dots and flicker stimuli were corrected for multiple comparisons using the Bonferroni method at a significance level of 0.05. This analysis found 20 brain regions (volumes of interest (VOIs)) that responded to the moving dots stimuli. Out of these, 14 (70%) showed increased activity, and 6 (30%) showed decreased activity. For the flicker stimuli, 8 VOIs were identified, with 6 (75%) showing increased activity and 2 (25%) showing decreased activity. All significant brain regions are presented in Fig. 3 and specified in Table S2 (supplementary material). In this work, only the positive changing signal VOIs were addressed, as the negative regions of the moving dots stimuli used do not contain relevant information (Duarte et al., 2023). All positive signal clusters for the moving dots were merged for the subsequent deconvolution-GLM analysis, and the same procedure was repeated for the flicker stimulus.

**Fig. 3.**
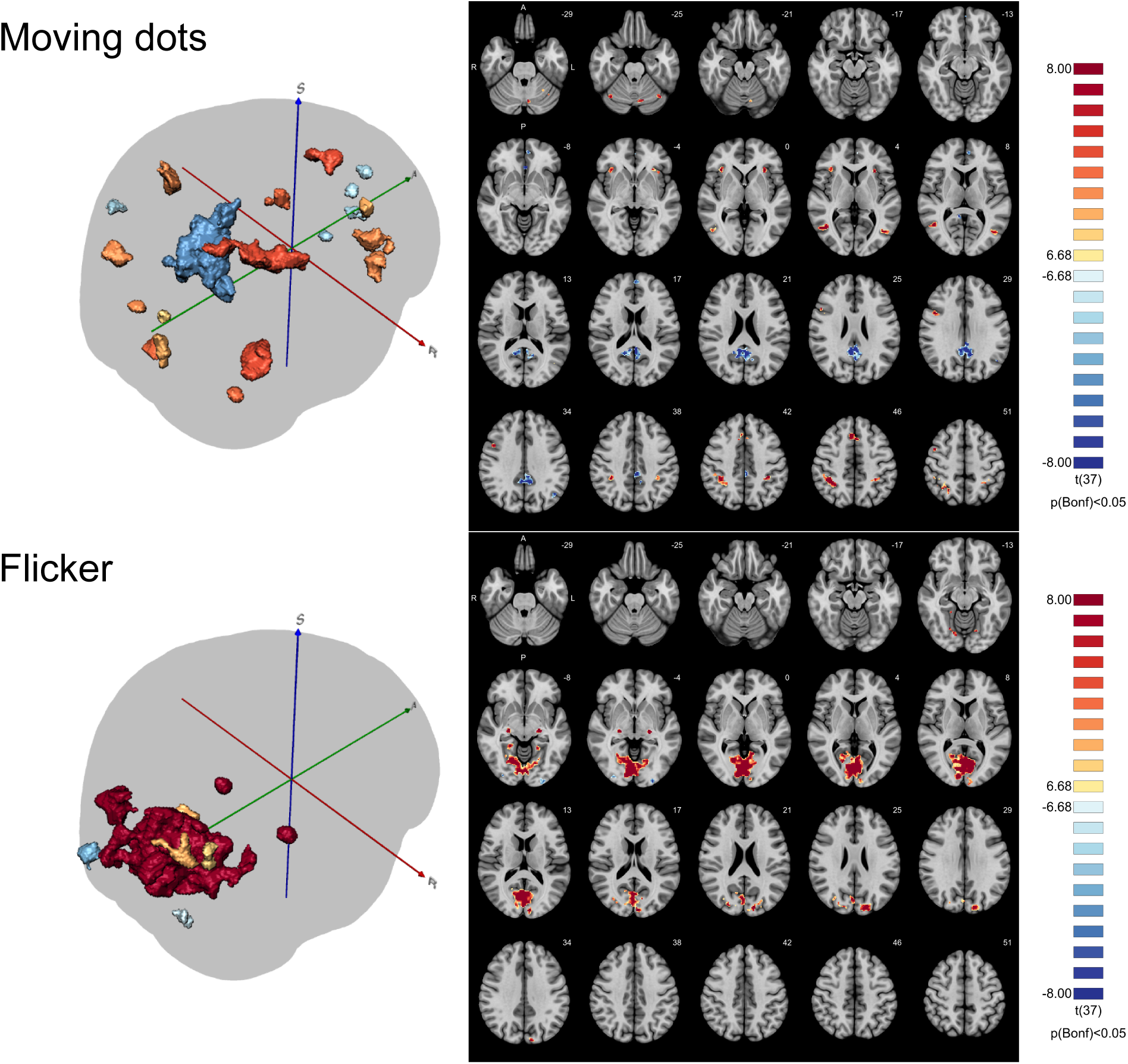
Functional maps generated from RFX GLM analysis of the fMRI response to any stimulation condition during the block design experiment for controls and type 1 diabetes mellitus patients. Moving dots left: Tridimensional representation of the brain activation map for the moving dots stimuli. Moving dots right: Transversal planes of the brain activation map for the moving dots stimuli. Flicker left: Tridimensional representation of the brain activation map for the flicker stimuli. Flicker right: Transversal planes of the brain activation map for the flicker stimuli. The maps are corrected for multiple comparisons using Bonferroni correction (p < 0.05). One can see significant positive signal change (yellow/red) and negative signal change (blue) in several clusters. We used the positive signal changing VOIs to extract the fMRI BOLD signal for further analysis of the hemodynamic response function in each group. Transversal planes are shown in the right panes. S, superior; A, anterior; R, right, L, left, P, posterior.

### 2.13. Deconvolution of brain hemodynamic responses

The fMRI BOLD signals measure changes in oxygenated blood levels over time. However, this signal is noisy, and a single stimulus does not allow us to determine the HRF. In event-related designs employing brief stimuli, the HRF, typically lasting around 20 s (Logothetis and Wandell, 2004; Taylor et al., 2018), can be estimated using a deconvolution approach. Indeed, individual NVC responses overlap and combine in the recorded BOLD signal. Therefore, to separate these overlapping signals, a deconvolution-GLM is used. This method utilizes the Moore–Penrose pseudoinverse, along with the specific timing of stimuli, to estimate the impulse response function. Unlike the standard GLM, the deconvolution-GLM does not assume a predefined HRF shape; therefore, it produces beta weight estimates based solely on the data (Serences, 2004; David et al., 2008). Although some authors compute the HRF for each subject and the group HRF as the average or median of the individual HRFs, in this work, the HRF for each group was established considering all acquisitions simultaneously, as each constitutes a probe for the group. Additionally, event-related moving dot stimuli, featuring faster motion presented alternately on the left and right, were combined. The deconvolution-GLM was computed for all three stimulus conditions (Threshold, Sub-maximum, and Flicker) within the VOIs of positive BOLD signal change, separately for the healthy and T1DM groups.

### 2.14. Statistical analysis

The estimated brain hemodynamic responses were compared between groups using the t-test for independent samples at each time point, based on the mean and standard error of all the measurements of the BOLD signal for all subjects in each group. Resulting p-values were corrected for multiple comparisons using the Benjamini–Hochberg procedure. Time points with corrected p-values of less than 0.05 were considered statistically significant in showing differences between groups or subgroups.

## 3. Results

### 3.1. Hemodynamic response patterns in the brain

BOLD group responses were analyzed for healthy participants (n = 16) and individuals with T1DM (n = 20) across three visual stimulus conditions: a Threshold-level speed discrimination task, a Sub-maximum speed discrimination task, and a flickering light stimulus. Results are shown in Fig. 4. Interestingly, the T1DM group exhibited a trend toward higher BOLD responses across all three stimulus conditions. However, no statistically significant differences were observed between groups during the positive response phase (i.e., around the peak of the hemodynamic response). Both groups show a typical biphasic HRF: initial rise (positive peak) followed by a decline across stimulus conditions. Group differences appeared in the Sub-maximum and flicker stimulus in the undershoot region. Notably, for the flicker stimulus, group differences emerged at four distinct regions across the BOLD time course, specifically involving the initial dip and the post-peak undershoot. One of these regions, located within the undershoot phase, associated with the recovery to baseline, showed a significant group difference that persisted for over 25% of the total response duration. Puzzlingly, the T1DM group exhibited higher positive BOLD responses with slight temporal delays compared to healthy controls across all three stimulus types. For the speed discrimination tasks (both Threshold and Sub-maximum), the T1DM responses appeared as scaled-up versions of those in the healthy group, suggesting a potential exaggeration of the neurovascular response to motion stimuli. In contrast, markedly different response patterns emerged during the undershoot phase following the flicker stimulus, indicating stimulus-specific alterations in the post-activation vascular recovery dynamics.

**Fig. 4.**
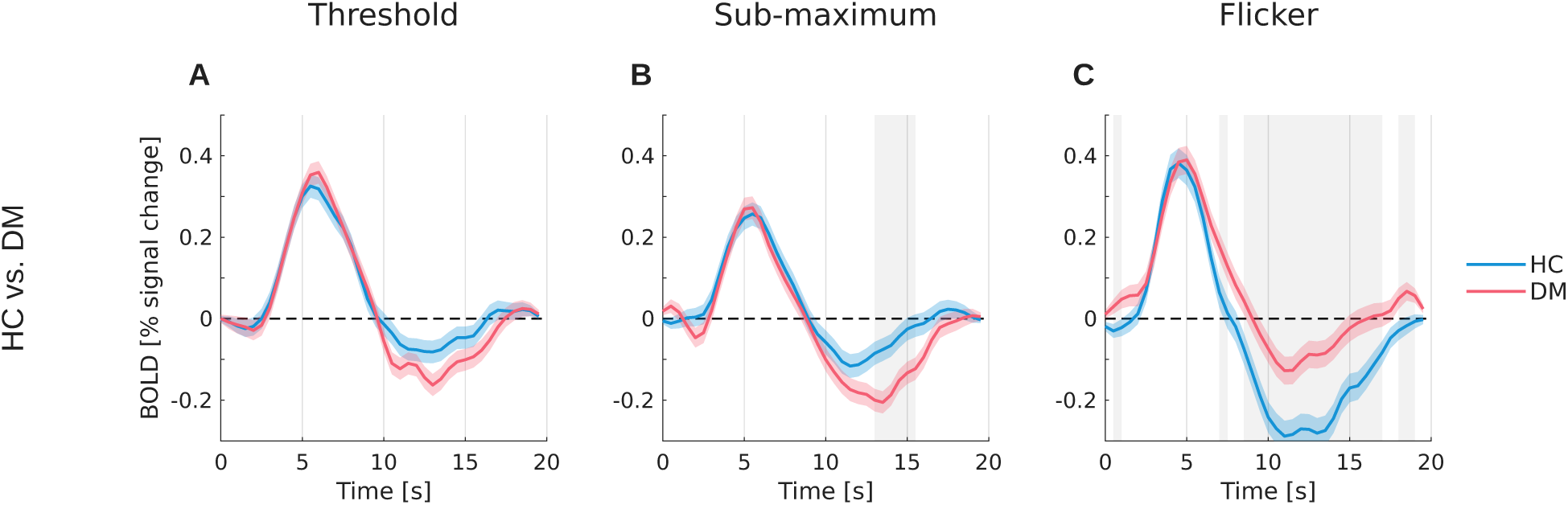
Hemodynamic response curves of the healthy and T1DM groups obtained using deconvolution-GLM. The solid lines represent the average beta weight with their standard error appearing in the surrounding region. The statistically significant analysis results (p < 0.05) of the differences between the two groups at each time point are represented in gray. Fully characterization of the p-values is presented in Fig. S3 (supplemental material).

### 3.2. Cortical hemodynamic responses in the (non-)responsive retina vascular network subgroups

Results associated with the cortical hemodynamic responses are presented in Fig. 5 with a simplified analysis in Table 1.

**Fig. 5.**
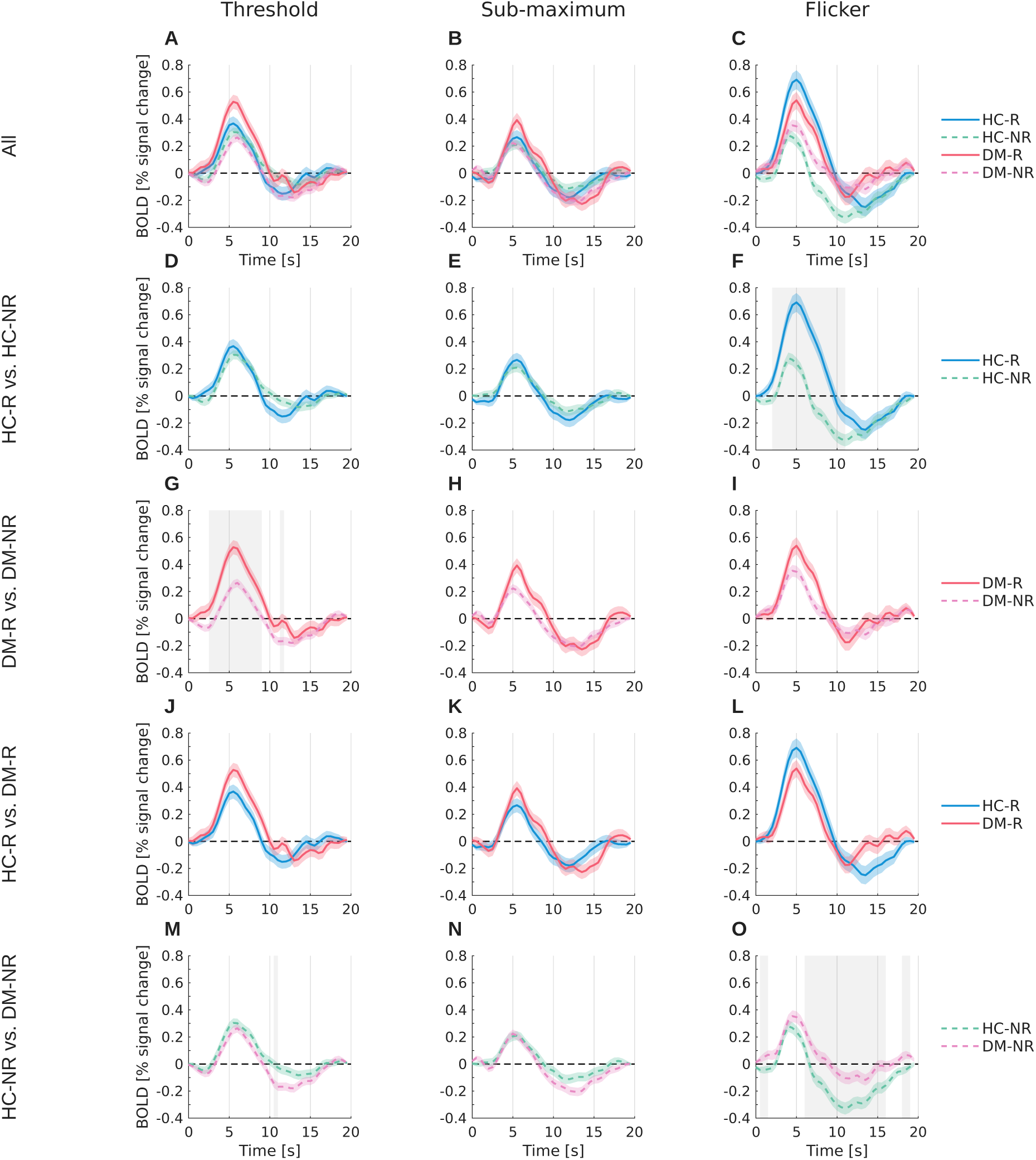
Hemodynamic response curves of the healthy and T1DM subgroups (with and without contralateral retinal response) obtained using deconvolution-GLM. The solid lines represent the average beta weight with their standard error appearing in the surrounding region. The statistically significant analysis results (p < 0.05) of the differences between the two groups at each time point are represented in gray. A to C: all four subgroups. D to O: paired subgroups with statistical comparison results. A simplified analysis is presented in Table 1.

Subgroups exhibiting responsive retinal vascular networks to contralateral stimulation — indicative of interocular effects (HC-R and DM-R, shown with solid lines) — displayed higher cortical BOLD signals compared to their non-responsive counterparts (HC-NR and DM-NR, shown with dashed lines) across all three stimulus conditions used in the fMRI experiments (Figs. 5, D to I). Statistically significant differences in the BOLD signal time course (gray-shaded regions) were observed only in two conditions: a) a stronger positive response to the flicker stimulus in the healthy subgroups (HC-R vs. HC-NR), with a minimum p-value at t = 7 s (p < 10⁻⁸; Fig. 5F), and b) a differential response to the Threshold-level speed discrimination stimulus in the T1DM subgroups (DM-R vs. DM-NR), with significant differences in the time interval 4.5 to 6 s (p < 0.001; Fig. 5G).

No statistically significant differences were observed between the responsive subgroups (HC-R vs. DM-R) across any of the three stimulus conditions (Figs. 5, J to L). In contrast, significant differences emerged between the non-responsive subgroups (HC-NR vs. DM-NR) during the Threshold condition (Fig. 5M) and flicker stimulus (Fig. 5O). Notable divergence in the BOLD signal time course of the flicker stimulus, particularly at the initial dip (immediately following stimulus onset) and during the undershoot phase (post-peak recovery), reaching a minimum p-value of < 0.003 within the 9–11 s interval. Moreover, HC-NR and DM-NR subgroups showed significant differences for the Threshold speed discrimination task, with a minimum p-value of 0.031 in the time interval 10.5 to 11 s. All statistical outcomes are detailed in Fig. S4 (supplementary material).

Finally, although no significant differences were observed between responsive T1DM and responsive healthy subjects (DM-R vs. HC-R; Figs. 5, J to L), the data in Figs. 5, G to I and 5, M to O reveal significant differences between non-responsive T1DM subjects and both their responsive counterparts (DM-NR vs. DM-R) and non-responsive healthy subjects (DM-NR vs. HC-NR).

## 4. Discussion

This study builds upon the novel findings on physiological crosstalk between human retinas reported by Jordão et al. (2025) to predict brain NVC responses in both healthy and T1DM subgroups, as measured by their BOLD signals. By employing various stimulation paradigms, we successfully characterized the brain’s hemodynamic response based on the presence or absence of a contralateral retinal response. Notably, the speed-discrimination task proved most effective in distinguishing between the two T1DM subgroups, whereas the flicker stimulus was more sensitive for differentiating healthy individuals and non-responder subgroups.

The utility of fMRI and its derived BOLD signal for investigating the human CNS is well established, despite ongoing debates regarding specific findings (Grinband et al., 2017; West et al., 2019) and challenges in data interpretation (Buxton et al., 2004; Kim and Ress, 2016). Nevertheless, this technique has been instrumental in advancing our understanding of NVC—a fundamental process essential for proper brain function and one that is significantly disrupted in numerous neurological and systemic diseases (Iadecola, 2017). Despite its importance, the precise mechanisms underlying NVC remain incompletely understood (Nippert et al., 2018). A recent study by Jordão et al. (2025) revealed a previously unrecognized NVC phenomenon in which photic stimulation of one retina can induce a vascular response in the contralateral retina, observed in a subset of healthy individuals and a comparable fraction of T1DM subjects without clinical signs of diabetic retinopathy. Building on this discovery, our findings demonstrate that stratifying subjects based on the presence or absence of a contralateral retinal vascular response reveals statistically significant differences in brain NVC between the subgroups within both healthy and T1DM populations. Furthermore, the findings presented here suggest that the absence of significant group differences reported in prior literature, particularly in studies on aging (Grinband et al., 2017) and non-diabetic diseases, may stem from an unrecognized heterogeneity within study populations, due to the uncontrolled mixing of physiologically distinct subgroups within both healthy and diseased cohorts.

Although distinct subgroups were identified within both the healthy and T1DM populations in this study, the comparable distribution of responsive and non-responsive individuals in each group, 5 out of 11 (45%) in the healthy group and 7 out of 16 (44%) in the T1DM group, supports the validity of direct comparisons between these populations. Based on this balanced stratification, we observed that T1DM subjects exhibited an overall higher NVC response in the brain across all three stimulus types. This finding contrasts with previous reports in type 2 diabetes mellitus (T2DM), where reduced BOLD responses relative to healthy controls have been consistently documented (Hu et al., 2019; Duarte et al., 2023).

Notably, greater statistically significant differences were observed in response to the flicker stimulus, which engages a broader spatial area and predominantly activates the visual cortex. In contrast, the Threshold and Sub-maximum stimuli elicited BOLD responses primarily in the inferior parietal gyrus (excluding the supramarginal and angular gyri) and the middle temporal gyrus, regions associated with attentional processing required for discriminating the relative speed of two moving points. Interestingly, in the positive response phase of the BOLD signal, the responses of T1DM subjects consistently fell within the range defined by the lower and upper bounds of the healthy subgroups, suggesting a preserved or intermediate NVC profile in this population.

The observation that T1DM subjects exhibit BOLD signal responses similar to those of healthy individuals, particularly in the positive response phase, which is commonly assessed using parameters such as peak amplitude and time-to-peak (Arichi et al., 2012; Duarte et al., 2023), suggests a pathophysiological trajectory in T1DM that is markedly distinct from that of T2DM. In contrast, studies on T2DM have consistently reported delayed time-to-peak and reduced BOLD signal amplitudes (Rangaprakash et al., 2021; Duarte et al., 2023), indicating impaired neurovascular function. These findings underscore the importance of distinguishing between diabetes subtypes when interpreting brain NVC responses.

An immediate potential clinical application of these findings is the ability to classify individual T1DM patients into specific subgroups based on their retinal vascular response, thereby enabling inference of their proximity to a healthy neurovascular profile using a non-invasive retinal imaging technique alone.

Unexpectedly, diabetic patients who lacked evidence of interocular coupling exhibited BOLD responses that closely resembled those of healthy controls. This finding initially contradicts the expectation that diabetic individuals, particularly those at risk for neurovascular dysfunction, would show impaired cortical responses. However, the absence of interocular coupling may reflect a disruption in binocular integration mechanisms, such as callosal transfer or binocular neuron modulation, rather than a global deficit in neurovascular coupling. As such, the cortical BOLD response in these patients may reflect a more simplified, monocularly driven activation pattern that mirrors the unimodal responses seen in healthy individuals under baseline conditions. Alternatively, early-stage diabetic patients may retain relatively preserved neurovascular coupling, with interocular integration being a more sensitive early marker of cortical involvement. These results suggest that interocular coupling itself, rather than the presence of diabetes, may serve as a key modulator of cortical BOLD amplitude in response to visual stimuli. Future studies should explore whether the absence of interocular coupling in diabetes reflects an early compensatory adaptation or a prodromal indicator of cortical dysfunction.

The primary limitation of this study is the relatively small sample size and the focus on a single disease population (T1DM). To draw definitive conclusions and assess the broader applicability of these findings, larger, multicenter studies are needed. Future research should include additional clinical populations to determine whether the observed neurovascular coupling patterns are specific to T1DM or extend to other conditions such as Alzheimer’s disease and related neurodegenerative disorders.

In conclusion, this study demonstrates that stratifying subjects based on the presence or absence of a retinal vascular response to contralateral photic stimulation reveals two distinct subgroups within both the healthy and T1DM populations, each exhibiting clear differences in brain BOLD signal responses. Notably, differences between T1DM and healthy subjects were confined to only one of these subgroups, suggesting that prior inconsistencies in the literature may stem from unrecognized heterogeneity within study populations. The identification of these previously uncharacterized subgroups calls for a reassessment of the current understanding of NVC, including the consideration of novel mechanisms. Collectively, these findings open new avenues for research and raise critical questions about the potential of NVC responses as non-invasive biomarkers for disease detection and stratification in diabetes, as well as whether the absence of interocular coupling in diabetes reflects an early compensatory adaptation or a prodromal indicator of cortical dysfunction.

## Glossary

NVC: Neurovascular coupling
BOLD fMRI: Blood-oxygen-level-dependent functional magnetic resonance imaging
CNS: Central nervous system
HRF: Hemodynamic response function
T1DM: Type 1 diabetes mellitus

## Acknowledgments

We are grateful to Sónia Afonso and Tânia Lopes for all fMRI imaging acquisitions.

## Author contributions

J.J.: Software, Formal analysis, Validation, Writing – first draft & review. P.G.: Formal analysis, Validation, Writing – review & editing. P.S.: Formal analysis, Validation, Writing – review & editing. J.F.: Subjects inclusion/exclusion criteria, Methodology, Writing – review. M.M.: Methodology, Writing – review & editing. D.C.D.: Writing – review & editing. MP: Subjects inclusion/exclusion criteria, Writing – review. M.C.-B.: Subjects inclusion/exclusion criteria, Methodology, Formal analysis, Investigation, Validation, Writing – review & editing, Funding acquisition, Resources. R.B.: Conceptualization, Project administration, Supervision, Subjects inclusion/exclusion criteria, Methodology, Formal analysis, Validation, Writing – review & editing, Funding acquisition, Resources. R.B. is the guarantor of this work and, as such, had full access to all the data in the study and takes responsibility for the integrity of the data and the accuracy of the data analysis.

