## Supplementary data for "The novel phenomenon of inter-retinal coupling predicts cortical neurovascular responses in health and disease"

**Supplementary Material**:

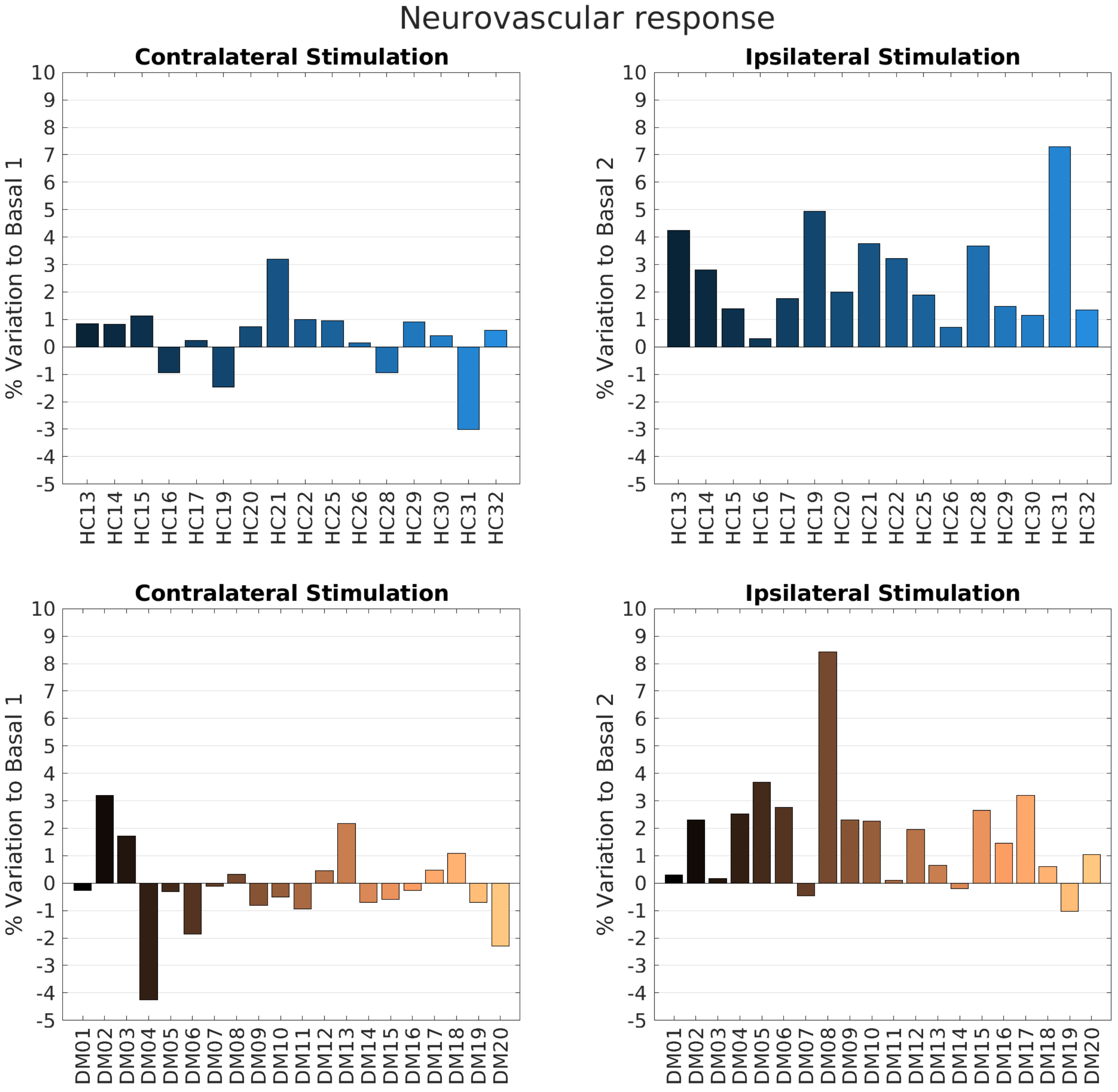

**Fig. S1. Neurovascular response data for the healthy control (top) and the T1 diabetic patient (bottom) groups.** Response to the photic stimuli is shown as a percentage of lumen width variation from the basal lumen width. Left: contralateral stimulation response. Right: ipsilateral stimulation response.

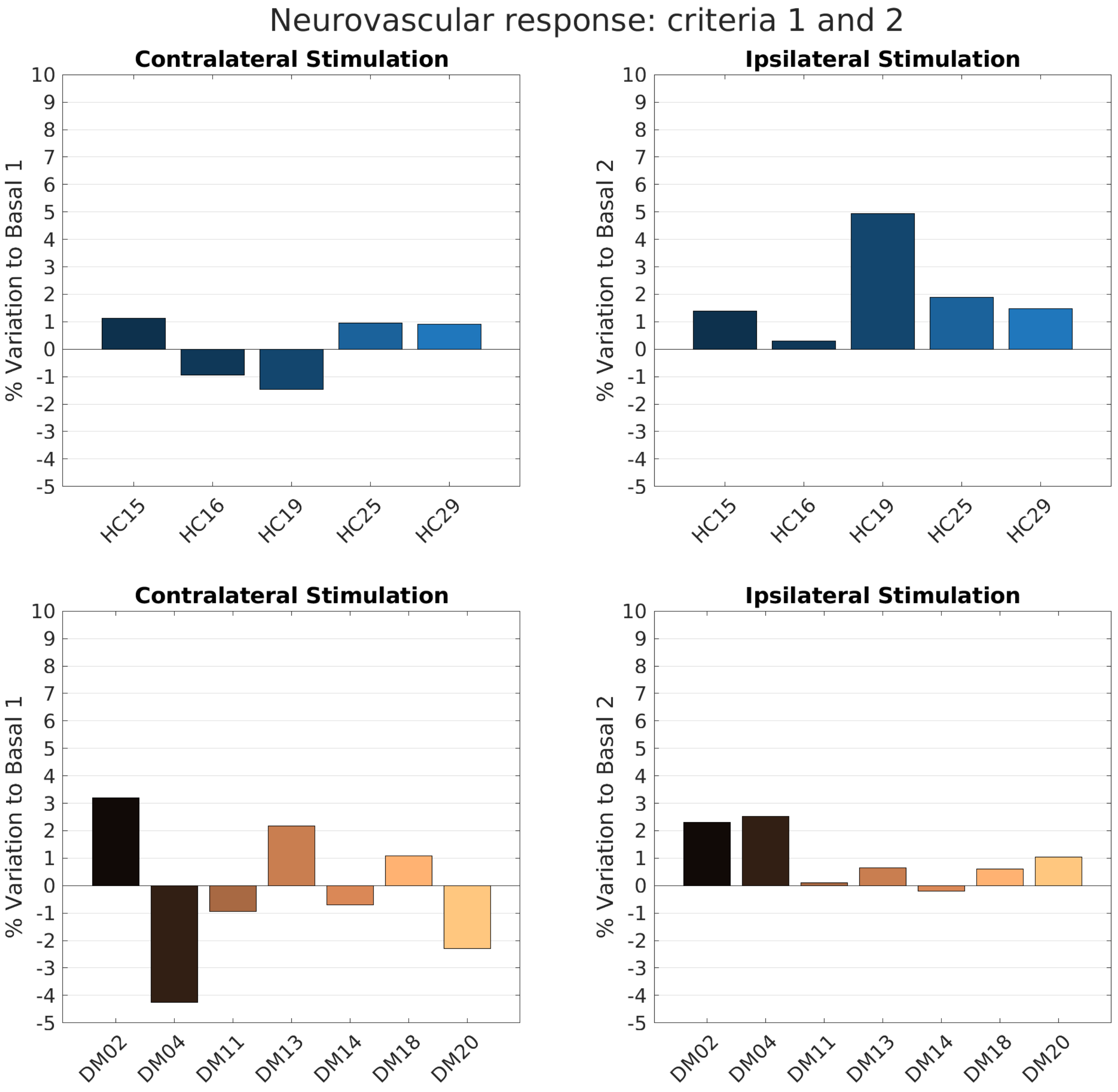

**Fig. S2. Neurovascular response data for the healthy control (top) and the T1 diabetic patient (bottom) groups after the use of selected criteria.** Healthy control cases (5/16 (31.3%)) (top) and type 1 diabetic patient cases (7/20 (35.0%)) (bottom) for whom the response to the contralateral stimulation recovered totally or partially to the basal lumen value and for which the response (absolute value) is over 1% or over half of the ipsilateral response for the same subject.

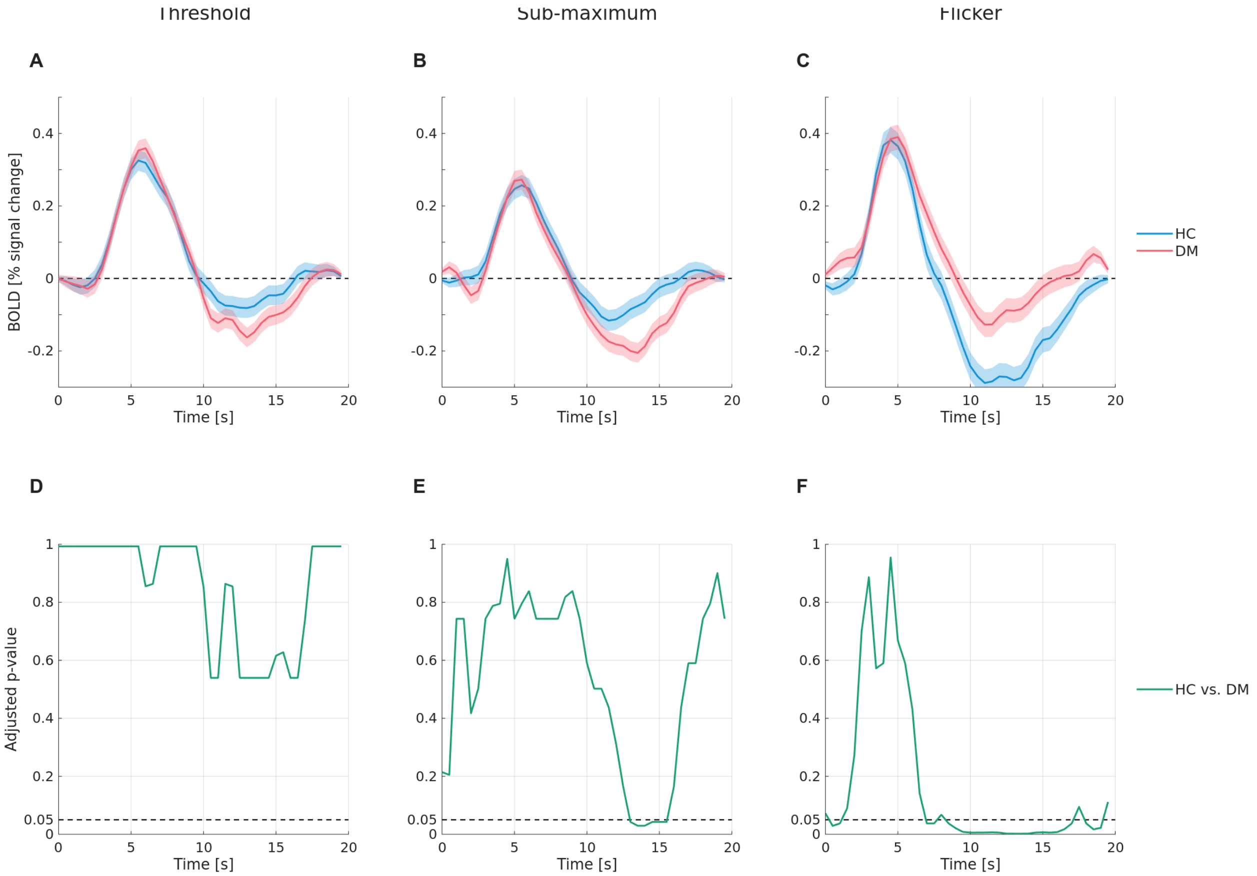


**Fig. S3. Hemodynamic response curves of the healthy and T1DM groups obtained using deconvolution-GLM with statistical analysis results.** Top row: hemodynamic response function (HRF) for the healthy control (HC) and type 1 diabetes mellitus (DM) groups for three stimuli: two speed discrimination tasks (Threshold and Sub-maximum), and Flicker stimulus. Bottom row: adjusted *p*-values for multiple comparisons. Only for the Threshold and Flicker stimuli (right column) a statistically significant difference between the two groups was found. Notably, the flicker condition significant differences extend for a large portion of the HRF, mostly at the post-positive-response undershoot region (panel F).

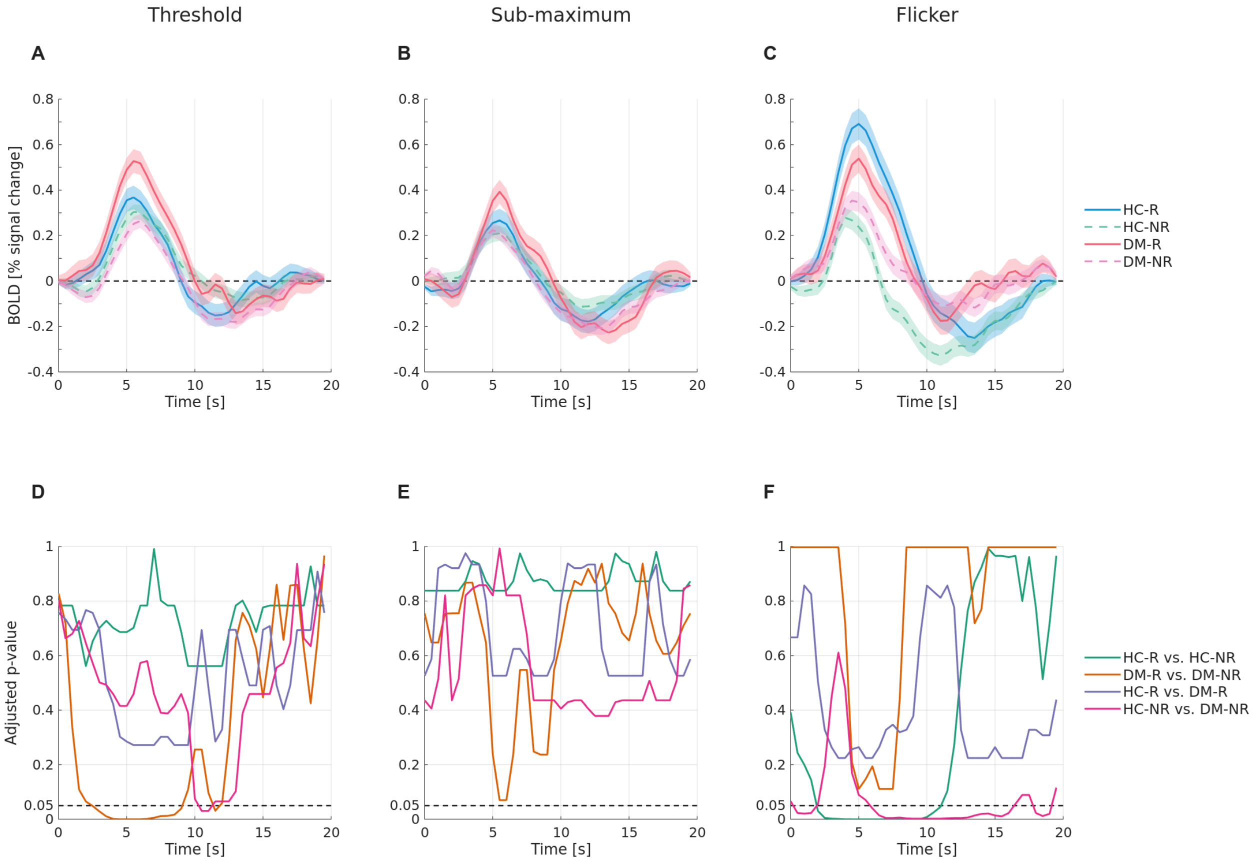


**Fig. S4. Hemodynamic response curves of the healthy and T1DM subgroups (with and without contralateral retinal response) obtained using deconvolution-GLM with statistical analysis results.** Top row: hemodynamic response function for the healthy control (HC) and type 1 diabetes mellitus (DM) responsive (R - solid lines) and non-responsive (NR - dashed lines) sub-groups (R/NR, vascular retina responsive/non-responsive to contralateral retina stimulation, respectively). Bottom row: adjusted *p*-values for multiple comparisons. For speed discrimination tasks, statistically significant differences were found only between the two sub-groups of the DM group at the positive-response region and between the non-responder groups in the undershoot region (panel D). On the other hand, the HRF present statistically significant differences between the two HC sub-groups in all but the raising edge of the positive-response region for the Flicker stimulus.

**Table S1. Raw data and subject characterization.** HC – healthy controls; T1DM – type 1 diabetes mellitus; M/F – male/female; F[M] – Menopause; OD/OS – right/left eye; BP – blood pressure; Spherical equivalent error (diopters) for the imaged (non-imaged) eye; IOP – intraocular pressure; Basal 1/2 – acquisitions without stimulation; Contra/Ipsi – acquisition after contralateral/ipsilateral stimulation; NA – not available.

| ID | Type | Sex | Age [years] | BP [mmHg] | HbA1c [%] | Spherical equivalent Error [D] | Imaged Eye | IOP [mmHg] | | Lumen width [pixels] | | | |
|---|---|---|---|---|---|---|---|---|---|---|---|---|---|
| | | | | | | | | Imaged eye | Contralateral eye | basal1 | contra | basal2 | ipsi |
| HC13 | HC | F | 19 | 122(59) | NA | -1.375(-0.875) | OS | 18 | 18 | 75.7 | 76.4 | 75.2 | 78.4 |
| HC14 | HC | M | 24 | 123(80) | NA | -0.25(-0.75) | OD | 16 | 16 | 96.6 | 97.4 | 95.7 | 98.3 |
| HC15 | HC | F | 24 | 110(80) | NA | -0.625(-0.5) | OS | 20 | 20 | 94.1 | 95.2 | 93.6 | 94.9 |
| HC16 | HC | M | 24 | 128(65) | NA | 0(0) | OD | 18 | 20 | 96.9 | 96 | 99.1 | 99.4 |
| HC17 | HC | F | 24 | 116(73) | NA | -0.875(-1.375) | OD | 23 | 23 | 76 | 76.2 | 76.6 | 77.9 |
| HC19 | HC | F | 24 | 116(73) | NA | -3.375(-2.5) | OD | 20 | 20 | 75.3 | 74.2 | 74.3 | 78 |
| HC20 | HC | F | 52 | 103(68) | NA | 2.5(2.25) | OD | 10 | 10 | 82.8 | 83.4 | 82.8 | 84.4 |
| HC21 | HC | F | 54 | 130(79) | NA | 0.75(0.875) | OS | 14 | 13 | 71.9 | 74.2 | 75.4 | 78.2 |
| HC22 | HC | M | 25 | 98(63) | NA | 0.25(0.5) | OD | 15 | 15 | 53.7 | 54.2 | 55.1 | 56.9 |
| HC25 | HC | F | 44 | 126(73) | NA | -2.5(-1.5) | OS | 12 | 12 | 63 | 63.6 | 64.4 | 64 |
| HC26 | HC | F | 45 | 102(65) | NA | -0.625(-1) | OD | 11 | 10 | 104 | 104.1 | 103.2 | 104 |
| HC28 | HC | F | 24 | 114(62) | NA | 1.375(4.375) | OS | 13 | 13 | 102.5 | 101.6 | 102 | 105.7 |
| HC29 | HC | M | 40 | 110(73) | NA | -4.25(-4) | OS | 11 | 11 | 64.5 | 65.1 | 65.1 | 66.1 |
| HC30 | HC | M | 55 | 110(70) | NA | -0.125(-0.25) | OS | 12 | 12 | 82.6 | 82.9 | 82.4 | 83.3 |
| HC31 | HC | M | 24 | 120(88) | NA | 0(0) | OS | 12 | 13 | 88.8 | 86.1 | 85.6 | 91.8 |
| HC32 | HC | M | 38 | 102(63) | NA | 0.875(0.25) | OD | 10 | 10 | 89.7 | 90.2 | 88.4 | 89.6 |
| DM01 | T1DM | F[M] | 45 | 90(66) | 6.7 | 0(0) | OD | 17 | 17 | 66.3 | 66.1 | 66 | 66.1 |
| DM02 | T1DM | M | 24 | 120(75) | 6.4 | -2.25(-2.75) | OS | 16 | 16 | 84.7 | 87.4 | 86.3 | 88.3 |
| DM03 | T1DM | M | 51 | 116(76) | 5.8 | -2.25(-2.25) | OD | 14 | 14 | 120.5 | 122.6 | 122.7 | 122.9 |
| DM04 | T1DM | M | 42 | 120(60) | 5.8 | -2.375(-2.5) | OS | 16 | 16 | 80.1 | 76.7 | 79.6 | 81.6 |
| DM05 | T1DM | M | 57 | 118(74) | 6.6 | 1.25(1.25) | OD | 14 | 14 | 78.7 | 78.4 | 79 | 81.9 |
| DM06 | T1DM | M | 57 | 127(79) | 7.4 | -1.125(-1.375) | OS | 16 | 14 | 72 | 70.7 | 70.5 | 72.5 |
| DM07 | T1DM | M | 22 | 122(74) | 6.7 | 0(0) | OD | 18 | 18 | 90.8 | 90.7 | 92 | 91.6 |
| DM08 | T1DM | F | 42 | 123(93) | 6.8 | 0(0) | OS | 12 | 12 | 72 | 72.3 | 70.1 | 76 |
| DM09 | T1DM | F | 28 | 96(71) | 6.9 | 0(0) | OD | 15 | 14 | 85.7 | 85 | 83.8 | 85.7 |
| DM10 | T1DM | F | 22 | 112(67) | 7.4 | -0.875(-0.375) | OS | 20 | 20 | 82.7 | 82.3 | 83.3 | 85.2 |
| DM11 | T1DM | F | 21 | 105(86) | 6.7 | 0(0) | OD | 20 | 20 | 111.7 | 110.7 | 112.3 | 112.5 |
| DM12 | T1DM | F | 47 | 121(73) | 7.9 | 0(0) | OD | 19 | 19 | 77.7 | 78 | 78.1 | 79.6 |
| DM13 | T1DM | M | 25 | 117(68) | 6.8 | -6(-6) | OS | 20 | 20 | 55.7 | 56.9 | 56.1 | 56.5 |
| DM14 | T1DM | M | 25 | 132(63) | 7.6 | 0(0) | OD | 21 | 21 | 82.9 | 82.4 | 82.7 | 82.5 |
| DM15 | T1DM | M | 27 | 139(72) | 6.8 | -0.5(-0.625) | OS | 16 | 16 | 103.4 | 102.8 | 103.9 | 106.7 |
| DM16 | T1DM | M | 44 | 136(86) | 7.6 | 0(0) | OD | 16 | 15 | 92.5 | 92.3 | 91.3 | 92.6 |
| DM17 | T1DM | F | 36 | 98(76) | 6.7 | -2.5(-3.5) | OS | 15 | 18 | 60.1 | 60.4 | 59.4 | 61.3 |
| DM18 | T1DM | F | 42 | 131(80) | 6.8 | 0(0) | OS | 14 | 14 | 82.9 | 83.8 | 83.3 | 83.8 |
| DM19 | T1DM | F | 20 | 113(68) | 7.3 | -1.25(-1) | OD | 21 | 20 | 108.6 | 107.8 | 107.2 | 106 |
| DM20 | T1DM | F | 24 | 103(69) | 7.2 | 0(0) | OS | 10 | 10 | 101.5 | 99.2 | 100.1 | 101.1 |

### Table S2. Brain clusters of statistically significant BOLD signal change with their peak MNI-space coordinates and respective AAL3 atlas labeling.

| Type | VOI | peak x | peak y | peak z | t | p | n voxels | AAL3 name | Hemisphere |
|---|---|---|---|---|---|---|---|---|---|
| **Positive signal** | | | | | | | | | |
| Moving Dots | 1 | 44 | -65 | 3 | 9.72 | 0 | 1007 | Middle temporal gyrus | Right |
| | 2 | 45 | 8 | 32 | 8.68 | 0 | 505 | Precentral gyrus | Right |
| | 3 | 38 | -44 | 46 | 11.00 | 0 | 2388 | Inferior parietal gyrus, excluding supramarginal | Right |
| | 4 | 44 | 1 | 52 | 7.85 | 0 | 141 | Middle frontal gyrus | Right |
| | 5 | 34 | 24 | -2 | 8.19 | 0 | 591 | Insula | Right |
| | 6 | 36 | -66 | -26 | 8.71 | 0 | 128 | Lobule VI of cerebellar hemisphere | Right |
| | 8 | 4 | 22 | 44 | 9.03 | 0 | 643 | Superior frontal gyrus, medial | Right |
| | 13 | -7 | -73 | -28 | 7.87 | 0 | 360 | Crus I of cerebellar hemisphere | Left |
| | 14 | -28 | -63 | -45 | 9.02 | 0 | 375 | Lobule VIII of cerebellar hemisphere | Left |
| | 15 | -37 | -41 | 41 | 8.25 | 0 | 712 | Inferior parietal gyrus, excluding supramarginal | Left |
| | 16 | -32 | 24 | 0 | 8.59 | 0 | 413 | Insula | Left |
| | 17 | -32 | -56 | -34 | 7.42 | 0 | 130 | Crus I of cerebellar hemisphere | Left |
| | 18 | -38 | -65 | -26 | 8.22 | 0 | 156 | Crus I of cerebellar hemisphere | Left |
| | 19 | -45 | -70 | 4 | 8.53 | 0 | 615 | Middle occipital gyrus | Left |
| Flicker | 2 | 28 | -73 | 24 | 7.34 | 0 | 184 | Middle occipital gyrus | Right |
| | 3 | -1 | -81 | 3 | 14.92 | 0 | 26726 | Lingual gyrus | Left |
| | 4 | 25 | -83 | 20 | 7.70 | 0 | 339 | Superior occipital gyrus | Right |
| | 5 | 22 | -22 | -6 | 10.87 | 0 | 256 | Lateral geniculate | Right |
| | 6 | -23 | -48 | -7 | 7.71 | 0 | 155 | Lingual gyrus | Left |
| | 7 | -24 | -22 | -8 | 11.68 | 0 | 319 | Lateral geniculate | Left |
| **Negative signal** | | | | | | | | | |
| Moving Dots | 7 | -3 | -48 | 28 | -9.69 | 0 | 7094 | Posterior cingulate gyrus | Left |
| | 9 | -3 | 55 | 17 | -7.69 | 0 | 177 | Superior frontal gyrus, medial | Left |
| | 10 | -4 | 28 | -9 | -8.16 | 0 | 101 | Anterior cingulate cortex, subgenual | Left |
| | 11 | -7 | 51 | -8 | -7.69 | 0 | 157 | Superior frontal gyrus, medial orbital | Left |
| | 12 | -7 | 50 | 7 | -7.54 | 0 | 118 | Superior frontal gyrus, medial | Left |
| | 20 | -47 | -66 | 33 | -7.79 | 0 | 217 | Angular gyrus | Left |
| Flicker | 1 | 31 | -91 | -5 | -7.24 | 0 | 134 | Inferior occipital gyrus | Right |
| | 8 | -30 | -99 | -6 | -8.74 | 0 | 337 | Undefined* (near) Inferior occipital gyrus | Left |
